# Neurons in inferior temporal cortex are sensitive to motion trajectory during degraded object recognition

**DOI:** 10.1101/2022.03.11.483956

**Authors:** Diana C. Burk, David L. Sheinberg

## Abstract

Our brains continuously acquire sensory information and make judgments even when visual information is limited. In some circumstances, an ambiguous object can be recognized from how it moves, such as an animal hopping or a plane flying overhead. Yet it remains unclear how movement is processed by brain areas involved in visual object recognition. Here we investigate whether inferior temporal cortex, an area traditionally known for shape processing, has access to motion information during degraded shape recognition. We developed a matching task that required monkeys to recognize moving shapes with variable levels of shape degradation. Neural recordings in area IT showed that, surprisingly, some IT neurons preferred blurry shapes over clear ones. Further, many of the neurons exhibited motion sensitivity at different times during the presentation of the blurry target. Population decoding analyses showed that motion pattern could be decoded from IT neuron pseudo-populations. Contrary to previous findings, these results suggest that neurons in IT can integrate visual motion and shape information, particularly when shape information is degraded, in a way that has been previously overlooked. Our results highlight the importance of using challenging multi-feature recognition tasks to understand the role of area IT in naturalistic visual object recognition. (Word count: 199)

## Introduction

Objects can be recognized even when visual information is ambiguous. For example, when walking through a foggy park, you might see a blurry shape sitting on the grass in the distance and be unable to figure out what it is. However, if the shape starts hopping, you may instantly recognize it as a rabbit. In this case, the stereotypical motion of the rabbit serves as a predictive cue for recognition. Here we present an investigation of the neural basis by which such predictive motion is used to recognize degraded visual images.

Past work has shown that behavioral demands can promote crosstalk between dorsal and ventral visual pathways, suggesting that motion and shape might not always be processed in isolation. As one moves along the ventral visual pathway, neurons exhibit selectivity for more complex features (Gallant et al. 1993; Rust & Dicarlo 2010). In early visual areas, neurons are sensitive to very basic object features, including orientation, size, luminance, contrast and motion (Hubel & Wiesel 1968; Schiller et al. 1976). Further along the visual pathway, V4 neurons are selective for simple combinations of these features, such as textures, curvature, and color (although the basis of their selectivity is still not well understood) (Touryan & Mazer 2015). Once visual information reaches inferior temporal cortex (area IT), the combinations of features become more complex in a way that is still not completely understood (Zhivago & Arun 2016; Rajalingham & DiCarlo 2019; Zhuang et al. 2021). However, it is well known that IT neurons respond to high order features (Tamura & Tanaka 2001; Op de Beeck et al. 2001) and that this area is implicated in representing categorical information for recognition (Logothetis & Sheinberg 1996; Kriegeskorte et al. 2008; Ritchie et al. 2015). It has been possible to identify 3D features that elicit maximal firing in IT neurons (Yamane et al. 2008) and to extract some computational relationships between features. But the semantic, real-world relationships between features that drive IT neurons remains an active area of research. Furthermore, the superior temporal sulcus (STS), which is anatomically situated between the dorsal and ventral visual pathways, is well known as a site of integration for shape and motion cues (Tanaka et al. 2002; Puce & Perrett 2003; Jastorff et al. 2012). However, much of the work in STS has focused largely on articulated motions (e.g. gestures, walking, motion of limbs), and little is known about the motion patterns of whole objects and their role in object recognition. Although area IT is bidirectionally connected to STS, there has been little evidence of motion processing in area IT, despite recent findings suggesting higher level representations of objects, including the physical properties of objects (Yildirim et al. 2019; Jia et al. 2021).

Moreover, previous behavioral tasks involving motion during object recognition have focused largely on other kinds of motion, including rotation (Vuong & Tarr 2004), structure from motion (Britten et al. 1992; Sáry et al. 1993; Beer et al. 2009), random dot motion (Pilly & Seitz 2009), articulated motion (Jastorff et al. 2006; Singer & Sheinberg 2010; Tyler & Grossman 2011; Schluessel et al. 2015), and complex naturalistic movies (Wang et al. 2012; Russ & Leopold 2015; Haxby et al. 2020; Jääskeläinen et al. 2021). While tasks using these kinds of motions can be informative, it remains largely unknown how trajectory motion plays a role in object recognition, even though trajectory motion is a common component of our perceptual experience. In some cases, motion can provide a strong cue to identification or even define the shape itself (Balas & Sinha 2008; Balas & Sinha 2009; Wang & Zhang 2010; Tian & Grill-Spector 2015). Thus, this experiment was designed to strike a balance between simple forms of motion, such as simple linear movements, and complex articulations that deform objects, to investigate the role of translational trajectories in neural processing during shape recognition.

To fully understand everyday vision, we must also understand processing of degraded visual information. Common circumstances we might encounter in daily life include recognizing objects in complex scenes among clutter, limited viewing angle, or poor lighting. Many methods have been explored to degrade visual stimuli in laboratory tasks, including decreasing contrast (Vogels & Biederman 2002; Namima et al. 2014), partial occlusion (O’Reilly et al. 2013; Tang et al. 2014; Namima & Pasupathy 2021), scrambling objects (Kravitz et al. 2010), morphing objects (Akrami et al. 2009), and salt-and pepper noise (Emadi & Esteky 2013; Kuboki et al. 2016). Most studies have shown that these methods characteristically reduce the magnitude of neural responses to visual stimuli both in V4 (Kosai et al. 2014), and in area IT (Nielsen et al. 2006; Emadi & Esteky 2013) during visual tasks. A few studies have shown that degradation can have mixed effects on neural responses in prefrontal cortex (Rainer & Miller 2000; Fyall et al. 2017) and that task experience can reduce degradation effects (Rainer & Miller 2000). However, it has recently been shown that some neurons in V4 can respond better to partially occluded shapes (Fyall et al. 2017), and that this is due to feedback from prefrontal cortex to area V4. The same group has also shown that neurons in V4 can respond with selectivity to blurred shapes, suggesting that degraded shapes are not always represented by decreased firing rates (Oleskiw et al. 2018). As area IT has a high degree of interconnectivity with area V4 (Felleman & Van Essen 1991), this suggests that *some* neurons in area IT might also respond better to degraded stimuli, particularly under behaviorally relevant circumstances.

Here we test the idea that reducing shape clarity will cause some neurons in area IT to respond to predictive motion during visual shape recognition. We designed a task that required monkeys to use motion information (2D motion trajectories) to recognize shapes under variable levels of shape degradation. The behavior of the monkeys on this task showed that monkeys can indeed use motion information to recognize degraded shapes. We simultaneously recorded from neurons in area IT while monkeys performed the active recognition task, as well as during passive viewing of the same moving stimuli. We hypothesized that neurons in IT that are selective for shapes are would also be selective for the corresponding, predictive motion trajectories. Given the role that predictive motion can play in recognizing degraded images, we further hypothesized that such tuning should be most readily revealed when viewing degraded images. Here we show evidence that IT neurons are selective for both shape and motion trajectories. The design of our experiments allowed us to further reveal that this shape-motion sensitivity is complex: shape selective neurons are not always selective for their predictive motion patterns, and motion sensitivity evolves over time at the population level. These findings provide an important first step in understanding the long-overlooked role of motion in the representation of shape within IT cortex.

## Materials & Methods

### Subjects & behavioral sessions

Two adult male macaque monkeys participated in the experiments. In both animals, access to area IT was made possible by use of a custom ball-and-socket chamber (Schiller & Koerner 1971; Sheinberg & Logothetis 1997) placed over either the right hemisphere (monkey Y) or the left hemisphere (monkey H) at +17mm anterior, +20mm lateral, (Horsley-Clark stereotactic coordinates). Chamber location was verified using CT scans for both animals, and the data was processed using a CT-MRI merge in NIH-AFNI software or the Brainsight system (Rogue Research) see **Figure 1A**. All surgeries were performed under isoflurane anesthesia, in accordance with the guidelines published in the National Institutes of Health Guide for the Care and Use of Laboratory Animals and approved by the Brown University Institutional Animal Care and Use Committee.

**Figure 1.**
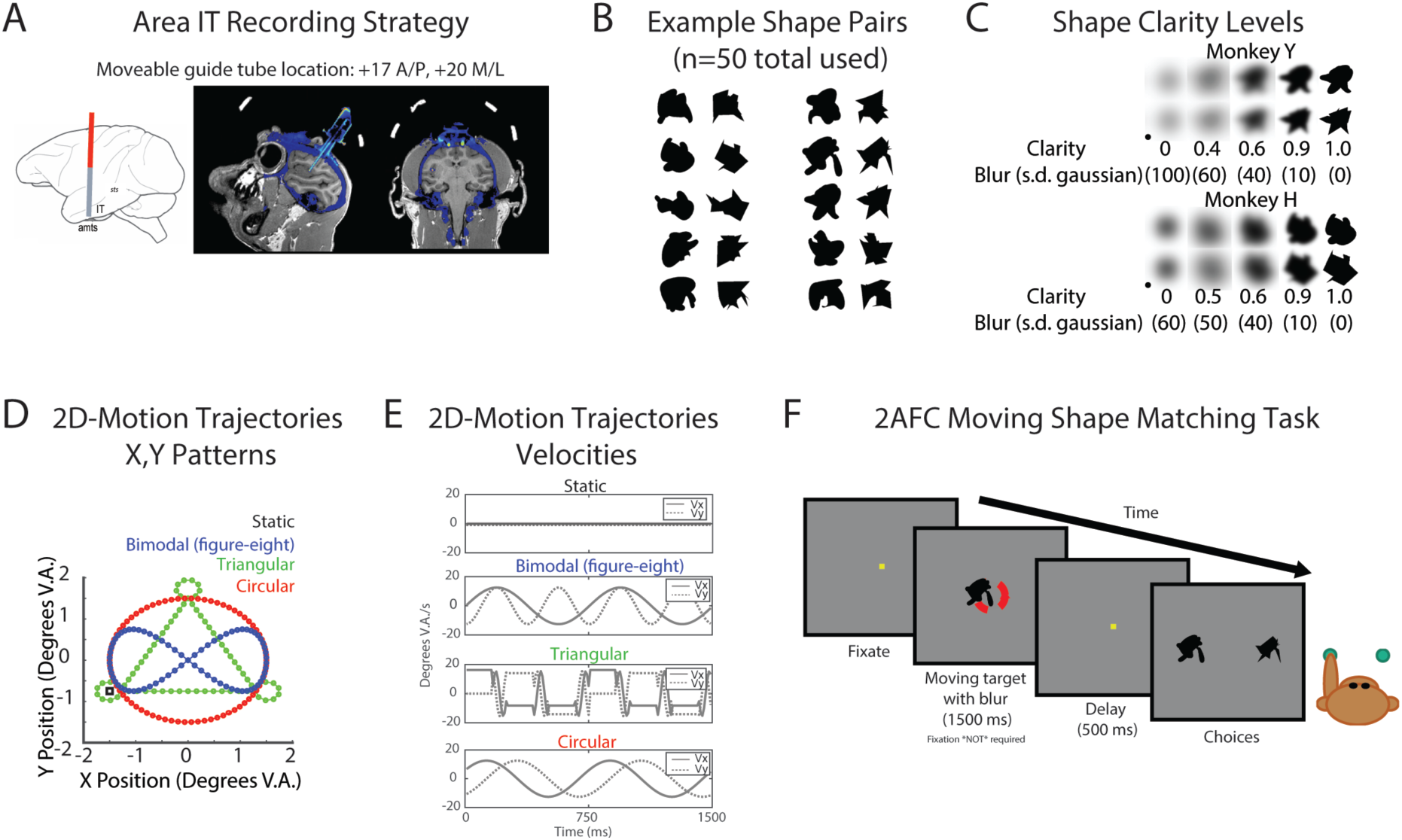
Task design and recording strategy overview. **A) Chamber placement and guide tube location for neural recordings from area IT**. Left: Sagittal atlas view of approximate recording approach Right: CT-MRI overlay for Monkey H, sagittal view (left), coronal view (right). **B) Example 2D shapes used in behavioral task**. 50 pairs of randomly generated 2D shapes were used in the behavioral task. Each pair, as shown here, consisted of one “blob” (first column) and one “spikey” (2nd column), and were generated from the same randomly overlapping polygons (n=4). 10 example pairs are shown. **C) Shape clarity levels used in behavioral task for each monkey**. Clarity (gaussian blur) levels are shown for both Monkey Y (top) and Monkey H (bottom). Shape stimuli were made more ambiguous by using a gaussian blur with a particular standard deviation (s.d.) in pixels. Clarity was defined as 1-(blur/100), except in the 0 clarity case. In this condition, for both monkeys, a black circle (see inset) was the underlying stimulus, such that there was no diagnostic shape information that could be extracted from the stimulus. **D) The four movement patterns used in the task**. The x,y positions of the four possible target motion patterns (Green=triangular, Red=circular, Blue=bimodal, Black square=static) are plotted. Stimuli started in the bottom left near the overlap of the three motion trajectories, except in the static case, where the location is marked with the square. **E) Target motion x, y velocity profiles**. These correspond to the motion trajectories in D. **F) Example trial of match-to-sample task with moving shapes**. The monkey first had to acquire fixation for ∼200 ms, after which the fixation spot disappeared and a moving target appeared for 1500 ms. The allowable eye position window (not shown) during the target period was -4 deg. V.A. in x and y to encourage pursuit of the moving target. After 1500 ms, the target disappeared and the fixation square reappeared for 500 ms for a delay period. At the end of the delay, the two static choices appeared and the monkey had 2 seconds to select an answer using a button press.

### Task stimuli

Two-dimensional (2D), randomly generated, geometrical filled shapes were used in the behavioral tasks. The shape set consisted of 50 pairs of shapes, or 100 shapes in total. Half of the shapes were sharp-edged polygons called “spikeys” and their smooth interpolated matches were called “blobs.” Example pairs are shown in **Figure 1B**. Shape stimuli were blurred by convolving the shape images with a circular 2D Gaussian kernel with variable standard deviation (s.d.) (Oleskiw et al. 2018). Five blur levels (i.e. clarity levels) were chosen based on each animal’s behavioral task performance, such that there were three levels between the two extremes of no blur (100% clarity) and 0% shape clarity (**Figure 1C)**. Shape clarity was defined as (1-s.d. of the 2D Gaussian)*100, except in the 0% shape clarity case, because the underlying shape was not a shape from the set. The underlying shape stimulus of the 0% clarity stimuli was a circle polygon (not the true shape) to ensure that there was absolutely no task-relevant shape information in the motion-only stimulus. Monkey Y had a fainter 0% clarity stimulus than Monkey H to reduce behavioral bias; the values of the s.d. of the Gaussian used to blur the circle are shown in Figure 1C. Three 2D motion trajectories were used in the task. The x,y position patterns of the three trajectories (circular trajectory, triangular trajectory, and bimodal (figure eight) trajectory) are shown in **Figure 1D**. The periodic trajectories were designed to last 1500 ms and have a 750 ms period (2 cycles total) to maintain similar average speed. Velocity profiles for the trajectories are shown in **Figure 1E**.

### Active match-to-sample task

In the two-alternative forced-choice (2AFC) task, the monkeys were required to report with a button press which of two choices best matched a target stimulus. On each trial, the monkey was first required to acquire the fixation spot (∼200 ms) within 1 deg. V.A. of the center of the screen, after which a target shape (3 deg. V.A.) appeared for 1500 ms duration. At target onset, the fixation spot was extinguished, and the fixation window was expanded to +/-4 deg. V.A. to encourage pursuit of the target. The target either moved in one of three ways, or not at all (static condition). After 1500 ms, the target was extinguished and the monkey was required to reacquire the fixation spot during the delay. After the delay (500 ms), two choices appeared. The monkey then had 2000 ms to press a right or left button to select which option of the shape pair best matched the target (**Figure 1F**). The shapes presented in the choice period were always 100% shape clarity and included the shape that matched the target shape and the corresponding blob or spikey shape. Trials with responses that were too slow (reaction times longer than 2000 ms) or absent were treated as aborts and were shuffled back in with remaining trials for repetition later within the block. Each shape category was associated with a motion pattern (Monkey Y: blobs-circular, spikey-triangular; Monkey H: blobs-bimodal, spikey-triangular). Both shape categories were associated with the remaining motion pattern and the static condition.

The task conditions were defined by the shape clarity, motion pattern, and shape category (blob or spikey). Trials with no shape information were rewarded randomly on 50% of completed trials, as there was no correct answer. Half of the trials for the session consisted of the single shape pair that was selected for most effectively driving the neuron(s) in that session. The other half of the trials consisted of any shape from the 100 shape set and were included to reduce behavioral bias. Monkey Y completed approximately 5 blocks of 180 trials per session and Monkey H completed approximately 5 blocks of 126 trials per session. During training, monkeys were exposed to the degraded stimuli and difficulty was increased over the course of months by decreasing shape clarity and increasing the number of task conditions. Each monkey was first trained on the match-to-sample task with variable shape clarity and unmoving (static) shapes for one to two months. Then each monkey was trained on the predictive motions (Monkey Y: blobs-circular, spikey-triangular; Monkey H: blobs-bimodal, spikey-triangular) for multiple months before the shared motion pattern was added into the task. The monkeys then each had two to six months of experience with all the task conditions interleaved before neural recording sessions began. The behavioral data presented was collected after extensive training on all task conditions. In this dataset, all task conditions were randomly interleaved and counterbalanced for response side; blocks only varied in the number of repetitions for each monkey.

Behavioral performance was calculated as the proportion of correct trials out of the possible number of trials for a given task condition (motion, shape, and clarity). Average reaction times were calculated across all incorrect and correct trials for a given condition. Performance across sessions was calculated by averaging the performance across sessions for a given condition.

### Passive viewing tasks

Passive viewing tasks were used to assess shape selectivity and to select a stimulus pair that would drive neural responses during the active task. The three passive viewing tasks had (1) all the 100 shapes with no degradation (2) a single pair of shapes (blob, spikey) with multiple shape clarity levels and no motion, and (3) a shape-less blur moving with each of the motion patterns. During all passive viewing tasks, a fixation spot was shown on the screen and the monkey was required to maintain fixation on this spot for ∼250 ms before a stimulus appeared. Multiple stimuli (up to 9) were shown during a single trial, during which the monkey maintained central fixation. Each stimulus was repeated at least 10 times in random order. During passive viewing of the 50 shape pairs without motion, each shape was shown at the center of the screen (size approximately 3 deg. V.A.) for 200 ms with a 100 ms blank interval between stimuli. Online peristimulus time histograms were used to select a stimulus pair for the rest of the session. The shape pair used for a daily recording session was selected based on its effectiveness in driving the neuron recorded that day (or multiple neurons on an array) without any assessment for motion response. The single shape that evoked the largest firing rate from the neuron or majority of neurons was chosen and its corresponding match were used for the remainder of the session. During passive viewing of the selected single pair across clarity levels, each shape was shown for 200 ms (100 ms between stimuli) with all the clarity levels used during the match-to-sample task. During passive viewing of the moving blur (no shape information) the moving blur with one of the four motions was shown for the duration of the target in the active task (1500 ms).

### Eye movement data & analysis

Eye position was sampled by an IR based camera system (Eyelink) at 1kHz and moving averages were continuously stored to disk at 200Hz. At the beginning of each behavioral session, the eye position was calibrated to the computer monitor coordinates by running a standard nine-point calibration procedure (Stampe 1993). Average pursuit trajectories were calculated by averaging the x and y eye positions during the target period for each task condition.

### Neural data acquisition & analysis

Recordings in inferior temporal cortex (area IT, IT cortex) were conducted with single electrodes or 16-channel V-Probes (Plexon, Inc.). The neural data was amplified and digitized (25 kHz), and then processed in a Tucker-Davis Technologies (TDT) neurophysiology system. The raw data from each channel (electrode) was band-filtered with two frequency ranges: local field potential (LFP, 0.3-300Hz, downsampled to 1017Hz) and single units (SU, 300-3000Hz). Plexon offline software was used for manual spike sorting. Neurons were excluded from analysis if they were unstable over the course of a session or not driven by any visual stimulus or behavioral task feature and were regarded as task-irrelevant. Task relevant was defined as any significant modulation above pre-stimulus baseline (200 ms before stimulus onset) in five task time windows (typical IT visual: 50-200 ms, late visual 200-350 ms, very late visual 800-950 ms, early delay 1550-1700 ms, late delay: 1700-1850). It was clear that some neurons were not visual in the traditional sense, so we sought to remain agnostic to their role in the task without testing too many time windows (Welch’s *t*-test corrected for multiple comparisons α=0.008).

Sensitivity to shape clarity and motion were assessed for each neuron using either the early visual period (50-200 ms after stimulus onset) or a 150 ms sliding window (50 ms shift). Motion sensitivity at each clarity level was assessed with a one-way ANOVA for each shape class across motion patterns (α=0.05, corrected for multiple tests) across trials in each of the three motion conditions for each shape class. A traditional multifactor ANOVA was not tractable, as not all the motion and shape conditions were crossed: the “blobs” and “spikeys” each had a unique motion and only shared one motion pattern and the static condition. To assess motion sensitivity across the entire trial period, each unit was assessed as either being motion or shape selective using a one-way ANOVA using the same correction procedure for multiple tests.

Each session consisted of four phases: passive viewing of all the possible task shapes (static, i.e. not moving), passive viewing of a single shape pair across clarity levels (static), the active match-to-sample task, followed by passive viewing of the 0% shape clarity stimulus (4 motion conditions). Eye position, behavioral responses, and neural activity were recorded for all tasks.

### Population decoding analysis

Because neurons were recorded across sessions and not all simultaneously, a pseudo-population of neural responses to representative trials was constructed to represent the data across multiple sessions, serving as input into the decoding analyses. For each condition of interest, a subset of representative trials was randomly selected based on the maximum number of available trials across all sessions. For decoding of the four motion conditions, 64 randomly sampled trials (without replacement) were used for each clarity level, except for 0.0 clarity, where randomly sampled 96 trials were used for decoding. For each 150 ms time window, neural firing rates were concatenated as features for decoding each trial (# features = # of neural responses in the pseudo-population). The process was repeated for multiple iterations to capture more of the data in the decoding analysis and to reduce variability in decoding accuracy due to sampling bias.

Support vector machines (SVMs) were trained with a linear kernel using the MATLAB function fitcecoc.m from the MATLAB Statistics and Machine Learning Toolbox (MATLAB v2018b). Binary learners with ‘onevsone’ approach were used, meaning for each binary learner, one class was labeled as the positive class and another was labeled as the negative class; others were ignored. Cross validation within each step was done using crossval.m with a constant partition generated using cvpartition.m with kfold=# of trials, which resulted in leave-one-out training and testing of each trial repeatedly, across all trials in the single decoding analysis. Accuracy was computed using kfoldLoss.m of the cross-validated result, which produced a value of 0 or 1 for each trial indicating whether the individual trial was labeled correctly during the testing phase. Variance of the decoding accuracy was computed at this phase and then averaged across iterations (over re-selection of trials) to get the standard error of the mean decoding accuracy. A separate model was fit for each 150 ms time window (50 ms sliding window), following previous work demonstrating success in using these timing parameters for decoding neural responses in area IT (Babolhavaeji et al. 2014). Default parameters were used if not otherwise stated.

Shuffled label tests were performed to establish decoding performance at chance level to capture bias in the data. Previous work has shown that comparing performance to only theoretical chance level can lead to misinterpretation of decoder performance, as an increase in sample variance of data values will also increase the chance of rejecting the null hypothesis when compared against theoretical chance (Combrisson & Jerbi 2015). Thus, following other recent developments in decoding toolboxes (Bode et al. 2019), repeats of all original analyses using the original data (i.e. randomized trial selection, # iterations of a k-fold cross validation) were conducted using a random assignment of labels to exemplars to produce the “shuffled labels” level of chance. This accounted for any potential biases in the original data that might affect true chance level accuracy and would not be accounted for using theoretical chance levels (although both are shown in each decoding figure). The original and shuffled label analyses were otherwise identical.

## Results

We designed a task and recording strategy to explore the relationship between shape ambiguity and motion trajectory in area IT, an area known for shape processing. We hypothesized that when shape information is degraded, IT neurons responding to shape information would respond more to movement during recognition. Based on behavioral data, we show that the animals could recognize the motion trajectories and use them to select the correct shape even when shape was degraded. Analysis of IT neural responses during the task show that firing rates did not always decrease with shape degradation, and that neurons exhibited multiple kinds of responses to shape clarity. Moreover, neurons exhibited a variety of responses to stimulus motion in time. Analysis of motion sensitivity across the target and delay period and population decoding suggests that a significant proportion of neurons responded to motion throughout the target period. Lastly, we compared decoding performance from the active task (where smooth pursuit was allowed) to decoding performance from the passive task (fixation required), to demonstrate that decoding performance was not solely dependent on eye movements.

### Monkeys can learn to use motion to recognize degraded shapes

Two monkeys completed approximately 30 sessions of the match to sample task (33 sessions for Monkey H, 28 sessions for Monkey Y), which required matching a shape to a target that varied in motion and shape clarity. The easier shape clarity trials could be completed correctly using motion or shape information, whereas the difficult clarity levels, and particularly the condition with no shape information (0.0 clarity) required use of motion-shape associative knowledge. The motion trajectory mappings were different for the two monkeys, so the particular motion-shape associations were not critical as multiple mappings were learnable. As shape clarity increased, accuracy increased from chance level to around 90% for the (uninformative) motion conditions that both shape categories had in common (static and shared motion) (**Figure 2**, left). However, for the motion patterns uniquely associated with only one shape class (blob or spikey), performance was not affected by shape clarity and remained around 90% accurate across all shape clarity levels. This demonstrated that the monkeys were using the learned motion-shape associations to select correct responses, even when shape information was degraded or unavailable. Reaction times varied slightly across the motion conditions despite the 500 ms delay before choice period. Reaction times trended slightly faster (although not statistically different across every clarity level), for both monkeys in the conditions that involved use of the motion-shape associations (red & green curves, **Figure 2**, right).

**Figure 2.**
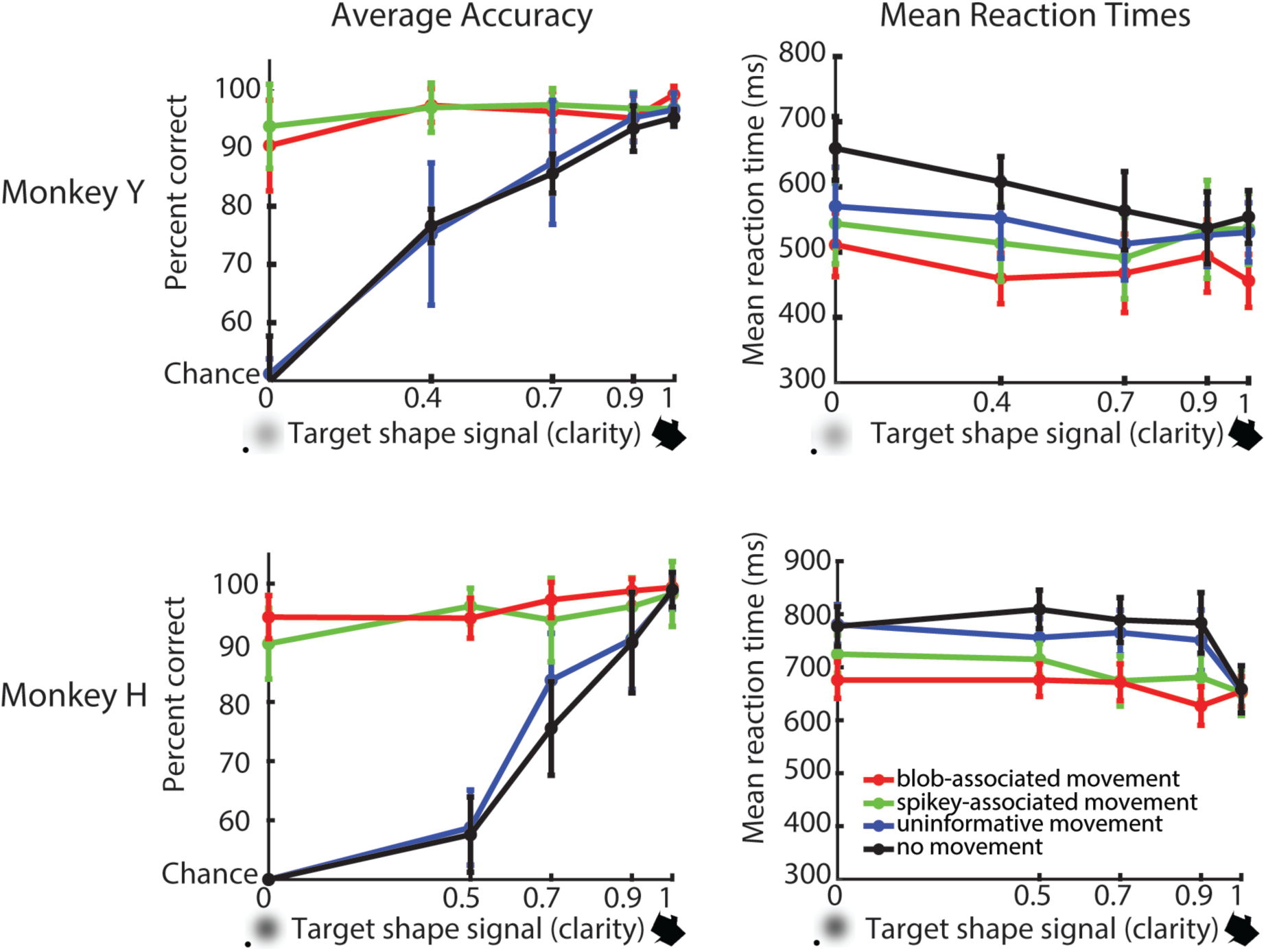
Behavioral performance and average reaction times during active task. Overall behavioral performance across sessions included in neural data analysis for both Monkey Y (28 sessions) and Monkey H (33 sessions). Average performance for each monkey (left) across the four motion conditions demonstrates the use of the motion-shape associations in low shape signal (clarity), as performance stayed near ceiling for the hardest clarity levels when the target was moving with the two predictive motions. Performance decreased to chance for the most difficult clarity level (target shape signal 0) for the uninformative motion and static conditions. Reaction times were slightly slower for the static condition for both monkeys (right, black curves). Error bars s.e.m. across sessions.

The goal was to create a behavioral task for monkeys to recognize and select shapes based on motion patterns. The behavioral data demonstrates that the monkeys can flexibly use motion information when shape information is not available to select the corresponding shapes. Both monkeys learned the motion patterns associated with each shape class to properly choose a shape that matched a degraded moving shape. We next asked how neural responses in area IT are affected by shape degradation and motion.

### IT neuron responses to shapes are modulated by shape clarity

During each behavioral session, activity from one or more neurons from area IT was recorded using single electrodes or multi-contact probes. Neurons were first assessed for responsiveness to each of the possible task shapes using passive viewing (n=100 shapes). The shape the elicited the highest firing rate (i.e. single blob or spikey) was selected from the set and this shape, along with its counterpart (blob or spikey), and these were used for the remainder of the session. Neurons were then assessed for sensitivity to shape clarity using the selected blob and spikey pair. The two shapes in the pair were shown with variable levels of shape clarity in a passive viewing paradigm. The pair was shown using at least the same clarity levels as those used in the active task. We had hypothesized that as shape clarity increased, firing rates would increase. Although some neurons exhibited increased firing rate with increasing shape clarity for both shapes (**Figure 3A, top**), we also observed two other kinds of responses: increased firing rate with shape clarity for one shape and not the other **(Figure 3A, middle)**, or maximum firing rate with intermediate shape clarity **(Figure 3A, bottom)**. We make no claims about the boundaries between these three categories as there was clearly a gradient of responses between these three examples and a strict classification would be somewhat artificial. Overall approximate distributions of the subpopulations suggest that many units in the population respond best to degraded shapes rather than clear ones (**Figure 3B**). These are not strict categories to classify neurons, rather, are only used to demonstrate the heterogeneity of responses to the degradation of shape information.

**Figure 3.**
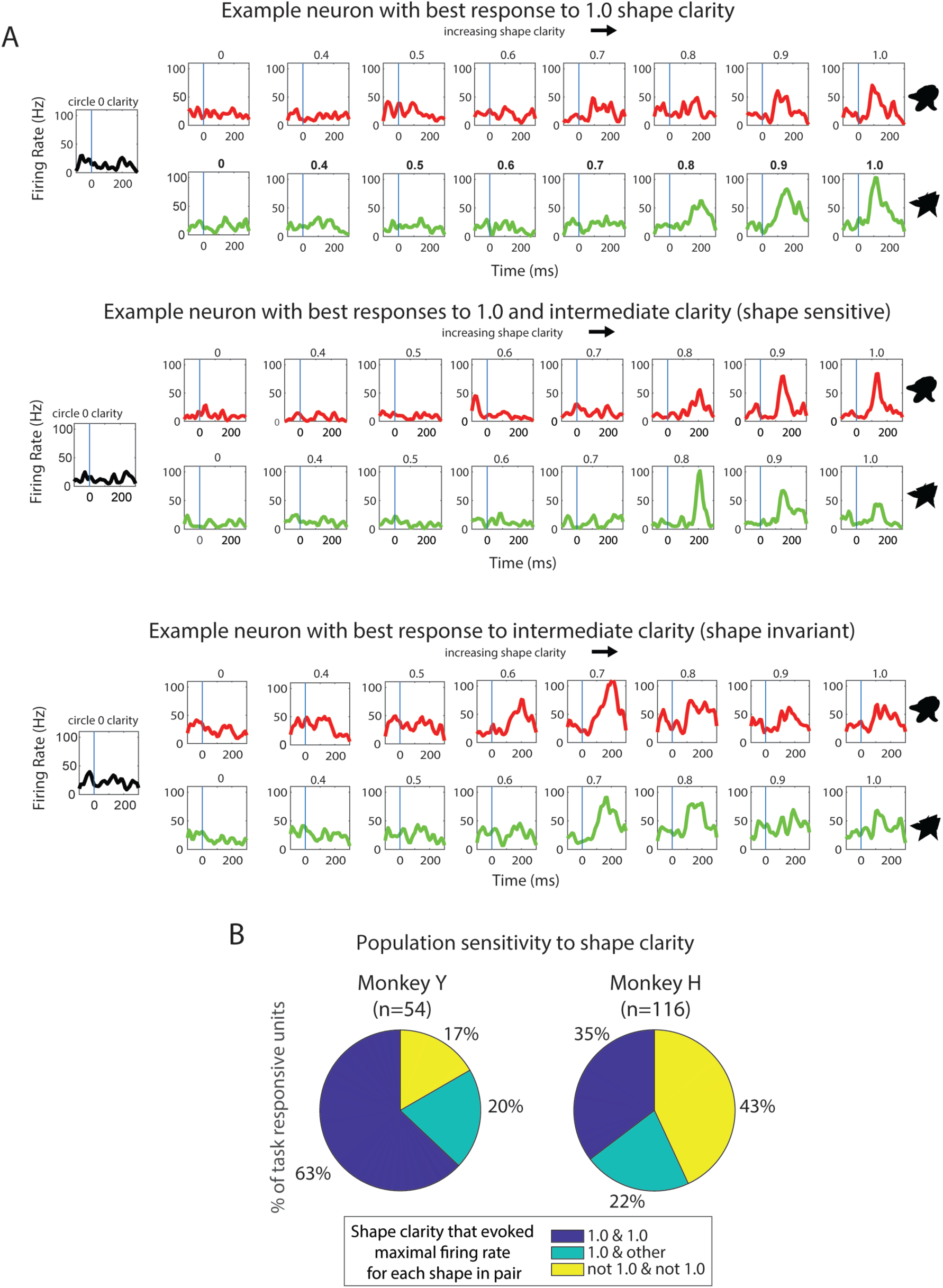
Multiple kinds of neural responses to clarity. **A) IT neurons can respond differently to varying shape clarity**. Top: An example of a neuron whose rate increases with increasing shape clarity for both shapes in a pair. Best response is at 1.0 shape clarity (no blur). Average firing rates for each shape (blob, red; spikey, green) at clarity levels ranging from 0 to 1.0, including all clarity levels in the task. Middle: An example of a neuron with a mixture of the kinds of responses seen in a) and b), that responds best to the blob with 1.0 clarity and spikey with 0.8 clarity. Bottom: An example of a neuron that responds best to 0.7 clarity for both shapes. **B) Quantification of the frequency of best responses to shape clarity**. Pie charts for each monkey show the distribution of neurons exhibiting the 3 kinds of responses described in (A). Left: Monkey Y, Right: Monkey H.

Given the observed variation of single unit responses to shape clarity, it was evident that traditional metrics of shape selectivity would not provide an accurate or useful representation of a neuron’s responsiveness to shape information. For example, neurons could sometimes exhibit their highest firing rate at 1.0 clarity for a blob, but an even higher firing rate for the corresponding spikey at lower clarity (**Figure 3B**). This type of response was not consistent across all shape pairs for this neuron, or for others, and thus not tractable with traditional metrics. Previously reported selectivity metrics, including degree of selectivity, breadth, and broadness (Mruczek & Sheinberg 2012) were calculated for all neurons, but no significant groupings or clusters were found, and there was no apparent relationship between the metrics and the three main kinds of responses observed. Thus, we chose to shift our approach to investigate how motion sensitivity, rather than selectivity, might change with shape clarity.

### IT neurons show clarity dependent sensitivity to motion

We asked whether neurons were motion sensitive and how motion sensitivity changed with target shape clarity. As neurons were not assessed for both shape clarity and motion with passive viewing, data from the active task was used for this analysis. To assess for motion sensitivity, firing rates from each motion condition for a particular shape stimulus (blob or spikey at a particular clarity) were compared using a one-way ANOVA (significance level *p*<0.05 then Bonferroni corrected for multiple comparisons). Thus, the main comparison was the neural response to a blob or spikey moving with the predictive movement pattern, the shared movement pattern, or no movement at all. The neurons exhibited a variety of responses to the motion conditions at various clarity levels (**Figure 4A)**. Surprisingly, there were multiple types of responses to motion of the same shape stimulus. Notably, motion sensitivity varied in time and occurred at a variety of clarity levels. The first subplot shows the neural responses (for unit Y2) to the predictive motion and shared motion at 0% clarity. In this example, the predictive motion evokes a significantly larger early visual response than the shared motion pattern for the same stimulus that has no relevant shape information. In contrast, the second subpanel shows an example of a unit for which the shared motion pattern evoked a much larger early visual response than the predictive motion pattern for a significantly blurred stimulus. Across this series of examples, the timing of and magnitude of peak activities in the single unit responses are modulated by motion type (shown by the colors in **Figure 4A**). A summary of the interaction between motion sensitivity and shape clarity during the early visual period is shown in **Figure 4B**. This plot shows the proportions of units that were motion sensitive at each clarity level 50-200 ms after target onset. Monkey Y had a larger proportion of units that were motion sensitive at intermediate clarity levels (but fewer units overall). There was a slight increase in motion sensitivity for both monkeys as clarity increased. Despite the differences between monkeys, this data shows that IT neurons can be sensitive to motion at different clarity levels. We next investigated motion sensitivity beyond the early visual period.

**Figure 4.**
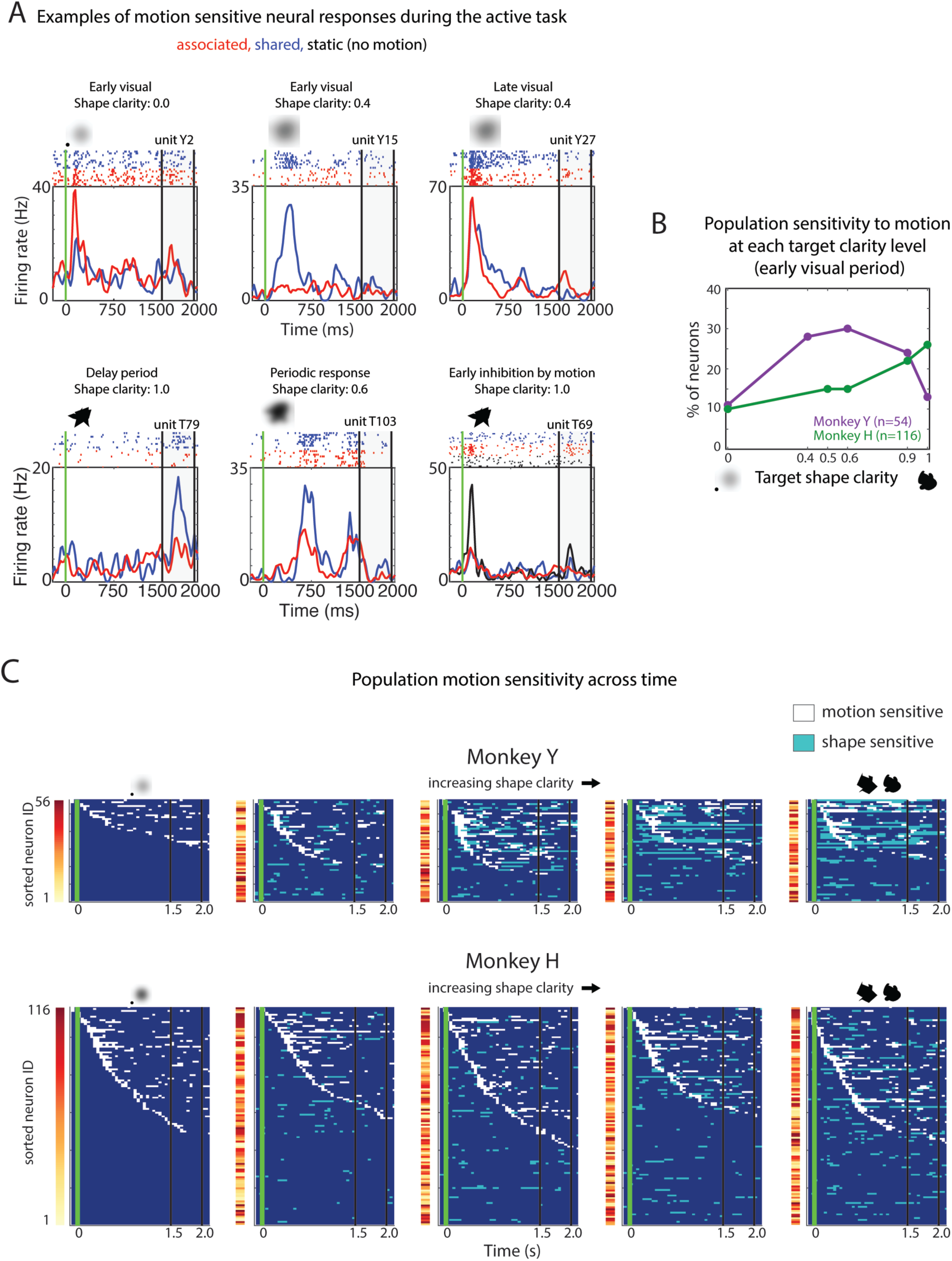
Multiple kinds of neural responses to motion and population tiling across time. **A) Single unit examples of motion sensitivity**. In each subplot, the response to the same shape stimulus (inset) to two different motion patterns (associated with that shape and not) is shown, revealing a variety of differences between motion conditions for the same shape. **B) Quantification of the proportion of motion sensitive units during the early visual period**. The plot is of the proportion of units that were motion sensitive at each target clarity, only during the early visual period (50-200 ms after stimulus onset). **C) Population motion sensitivity across the entire target and delay period for each clarity level**. In each subplot, the motion sensitivity (150 ms window, 50 ms sliding window) is shown across time. The neurons are sorted by which neuron had the earliest motion sensitivity for each subplot, revealing tiling across most of the target period and delay period for both monkeys (top—Monkey Y, bottom—Monkey H).

### Neural responses show dynamic motion sensitivity

The analysis above focused on the initial visual period, which is the time of typical visual responses in area IT. Previous work has suggested that degraded stimuli (such as the blurred ones used in this task) promote recurrent processing that might occur beyond the initial visual period (Wyatte et al. 2012; Tang et al. 2018; Ernst et al. 2021; Namima & Pasupathy 2021). Furthermore, the trajectory period was extended in time (1500 ms), thus it was natural to consider sensitivity to motion throughout the target period. We hypothesized that sensitivity to motion and shape information might fluctuate in time, and that the participation of units across the trial period might also fluctuate. To assess sensitivity to motion and shape information during the target period and delay, we used a sliding window analysis and sequential assessment of sensitivity to motion and shape class. Each unit was assessed as either being motion or shape selective (*p*<0.05, Bonferroni correction for multiple tests), representing significant modulation by either motion trajectory or shape class. **Figure 4C** shows motion sensitivity (white) and shape sensitivity (light blue) of individual units across the entire target and delay period at each clarity level, for both monkeys. The neurons are ordered by the time at which they are motion sensitive from earliest to latest. Critically, the ordering varies across clarity levels, shown by the color bar representing neuron identity to the left of each subplot. This suggests that the neurons that are motion sensitive at a particular time differ depending on the level of shape clarity. As shape clarity increases, the tiling of units decays for both monkeys, and appears not to extend as far into the delay period. Some units showed motion sensitivity at multiple time windows throughout the trial. No units showed sustained, significant motion sensitivity throughout the entire target period. This suggests that sensitivity to motion fluctuates in time and that the population could potentially represent motion patterns and support discrimination between motion conditions, even with degraded shape information.

### Population decoding shows decodable representation of motion information

Motion sensitivity analyses showed that different units were responsive to motion at different levels of clarity, but it remained unclear whether the pseudopopulations could discriminate the motion trajectories throughout the target or delay period. To determine if motion could be decoded throughout the trial, population decoding was carried out for the four motion conditions (static, shared, blob-predictive, spikey-predictive) for all clarity levels using SVMs (see *Methods: Population decoding analysis*). **Figure 5A** shows the average decoding performance for decoding 4 motion patterns for both monkeys for each clarity level. The decoding performance curves have a generally double-peaked shape in time, cycling around the time of the two cycles of motion (with some blurring from the smoothing and window size). Decoding accuracy is lower for 0.0 clarity yet above both theoretical chance and shuffled label decoding performance. To exclude the possibility that the decoders were mostly extracting information related to motion vs. static conditions, we also performed decoding on only the three movement trajectories and found that decoding performance remained above both theoretical and shuffled chance levels (**Figure 5B)**. As there is no object-related information present in these trials, the only way a decoder could extract information would be from motion information related to neural activity. Thus, these decoding analyses demonstrate that information about motion condition could be decoded from the pseudopopulations of area IT neurons. Because the monkeys’ eye movements were not constrained during the target presentation phase (i.e. they were allowed to pursue the shapes as they moved along the motion trajectories), we investigated if the decoding performance could be attributed to changes in eye position as opposed to object motion.

**Figure 5.**
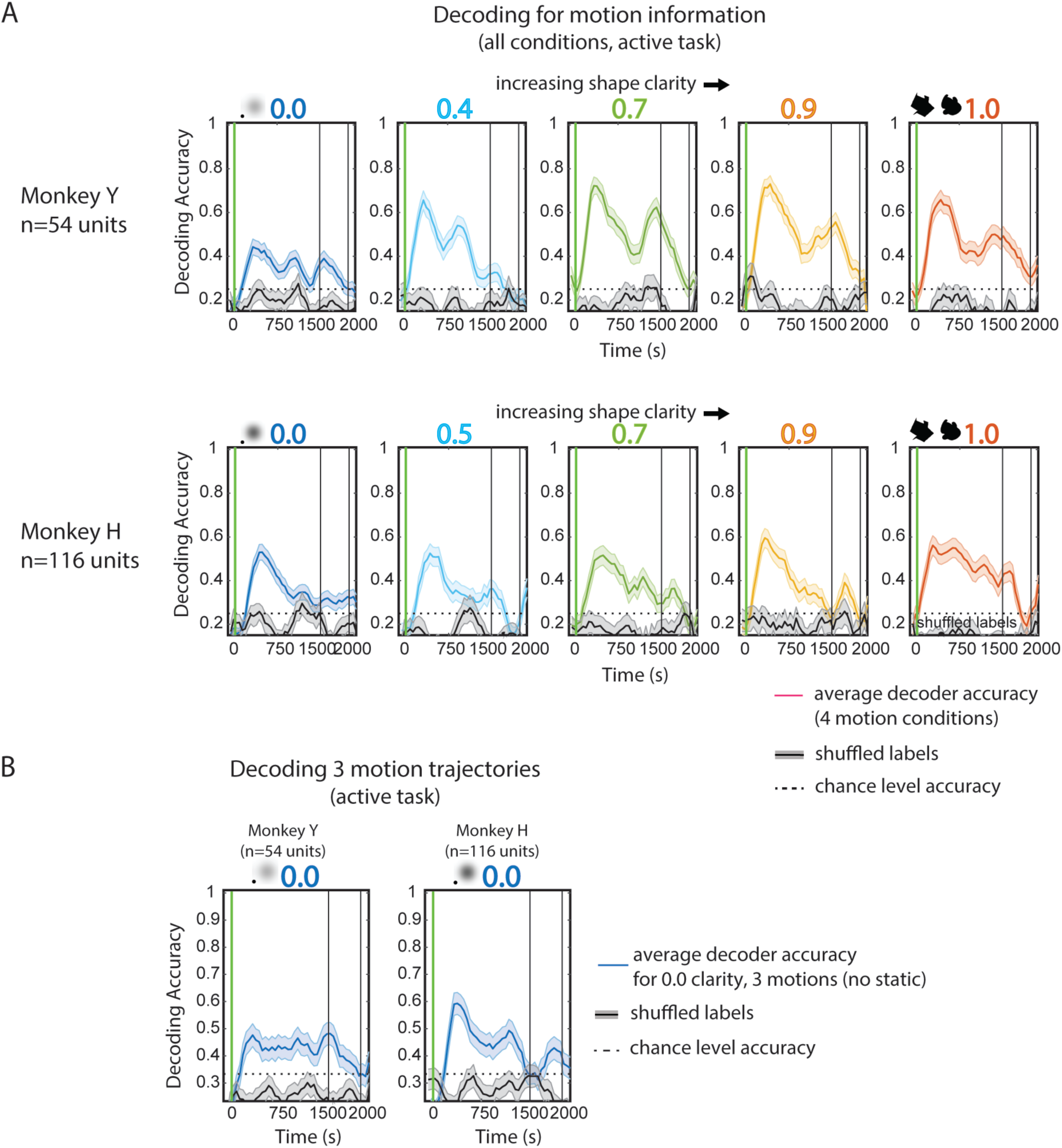
Decoding motion conditions from neural population activity. **A) Decoding for motion condition across clarity levels**. Multi-class (four-way) decoding of motion condition (static, shared, blob-predictive, spikey-predictive) was done using 150 ms windows, 50 ms sliding window; a separate model was used for each time window. Each panel is the average performance across iterations (of random resampling across trials and sessions) to decode 64 trials of combined neural activity across the ∼30 recording sessions. 0.0 clarity panels (far left) used 96 trials for decoding. Error bars s.e.m. Trial labels were shuffled and decoding analysis was repeated with the same parameters and numbers of iterations to create statistical chance level accuracy. Top: Average decoding performance for clarity levels 0.0 through 1.0 for Monkey Y. Bottom: Monkey H. **B) Decoding for motion trajectory without static condition at 0.0 clarity**. Multi-class (three-way) decoding of motion condition (shared, blob-associated, spikey-associated) was done using the same procedure as in (A), except using 72 trials for decoding. Error bars are the s.e.m. across iterations of decoding.

### Pursuit eye movements do not account for accuracy of motion decoding

During naturalistic object recognition, eyes naturally follow a moving object of interest. To allow for more naturalistic recognition in this matching task, the monkeys were not required to fixate. Rather, they were encouraged to pursue the moving targets with a large fixation window that required them to stay within +/-4 deg. V.A. of the center of the screen. Eye movement data demonstrated that both monkeys pursued the targets, and a transform of the original trajectory patterns were visible in the average eye movement patterns (**Figure 6A**). Average eye position data show anticipatory rightward saccades near the end of the target period (red line rightward) for both monkeys, reflecting a bias to acquire the right choice object first during the choice period. Both monkeys show some drift in eye position during the static conditions. Overall, the eye movement patterns were stereotyped and did appear to cover a smaller spatial range as clarity decreased (i.e. smaller eye movements were made). The eye movement data show that although the motion trajectory was the same across clarity levels, the eye movements in response to the motion and clarity changed across conditions.

Because the monkeys were pursuing the moving stimuli but not the static stimuli, it seemed possible that the population decoders were extracting information related to whether the monkeys were moving their eyes throughout the target period. However, as previously shown in Figure 5B, excluding the static condition from decoding analyses did not diminish decoding accuracy to below shuffled chance or theoretical chance levels. To further investigate the role of eye movements on the population decoding results, we performed decoding of the same three motion patterns during passive viewing, when the monkeys were required to maintain fixation (**Figure 6B**). Decoding of neural data during passive viewing was also significantly greater than shuffled and theoretical chance, supporting the idea that eye movements were not the sole contributor to decoding of motion condition in previously shown decoding analyses. Interestingly, decoding of data from the passive task showed a much more distinct bimodal shape than decoding of the neural data from the active task. This suggests that the decoders were likely relying on population activity related to other features than eye movements. However, it is possible that the small differences in retinal position of the stimulus across the conditions (perhaps due to quality of pursuit) could contribute to the decoding performance of motion patterns. This would imply that these IT neurons would have very refined spatial selectivity which was not otherwise apparent in the data.

**Figure 6.**
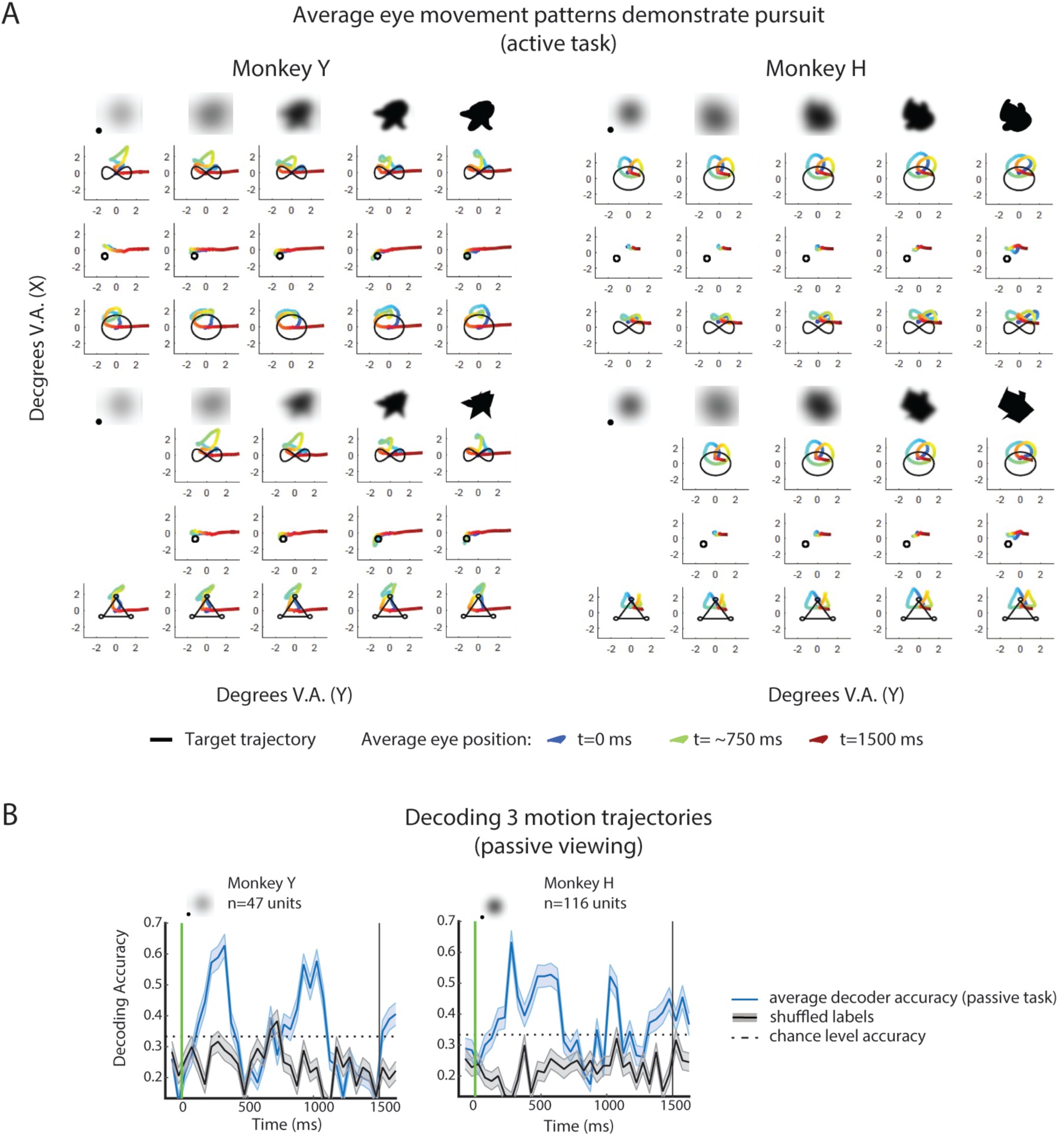
Eye movement data and decoding of motion-only conditions in active and passive tasks. **(A)Eye movement data demonstrate that both monkeys pursued the moving targets**. Average eye positions during the target period (1500 ms, color) are overlaid with the actual shape trajectories (black) across all shape + motion + clarity task conditions. Top rows show average eye movements for blob shapes across the three motion conditions (bimodal, static, predictive motion). Bottom rows show average eye movements for spikey shapes across all three motion conditions (bimodal, static, predictive motion). Average eye position is color coded from early to late (blue, green, red). Left: Monkey Y, Right: Monkey H. Averages are across all correct trials and behavioral sessions. **B) Multi-class (three-way) decoding of motion condition during the passive viewing task**. (shared, blob-predictive, spikey-predictive) at 0.0 clarity for both monkeys, using 36 trials for decoding. Error bars are s.e.m across iterations.

## Discussion

Our central finding is that neurons in area IT are sensitive to 2D trajectory motion in a task context where shape information is degraded or absent and motion can be informative. Firing rate analyses demonstrated that individual neuron responses to shape clarity and motion fluctuate in time. SVMs were used to decode activity of the pseudo-populations recorded from both monkeys to determine the extent to which the populations carried shape and motion information across different levels of shape clarity. Motion information appeared to be available in most clarity conditions, although monkey behavior suggested that the monkeys were using motion information primarily when shape information was not available, at lower shape clarity levels. Eye movements were permitted during the task to make the recognition of moving objects more naturalistic. Allowing pursuit does add an additional confound in our interpretation of the decoding results. However, decoding motion from the passive viewing of motion showed that eye movements were not solely responsible for the decoding of the active task. These analyses suggest that the population does indeed carry information about the motion trajectories.

The goal of the study was to investigate motion processing in area IT during moving object recognition. We began this study with the hypothesis that shape and motion are easily and naturally combined as visual features, and that because IT cortex plays such a critical role in object recognition, that it could have access to motion information under certain conditions. Specifically, we hypothesized that when shape information is degraded, informative motion information might support object recognition, and that neurons in IT cortex might be driven by motion information to compensate for a lack of shape information. As outlined in the introduction, there are multiple reasons why one might think IT neurons could play a role in the processing of motion during visual object recognition: the demonstration that motion can afford shape recognition (Marr & Nishihara 1978; Lawson et al. 1994; Mitsumatsu & Yokosawa 2003; Vuong & Tarr 2004; Friedman et al. 2009; Pike et al. 2010; Setti & Newell 2010; Nankoo et al. 2017), the role of IT cortex in coding temporal and structural information about visual associations (Sakai & Miyashita 1991; Morita & Suemitsu 2002; Eifuku et al. 2010), sequences (Meyer & Olson 2011; Ramachandran et al. 2017; Kumar et al. 2017; Kaposvari et al. 2018) and action patterns (Singer & Sheinberg 2010; Vangeneugden et al. 2011), and neural modulation based on behavioral experience (Li et al. 1993; Grill-Spector et al. 2006; Mruczek & Sheinberg 2007; Li & DiCarlo 2008). Furthermore, past work has also shown that prefrontal cortex carries motion and direction information (Zaksas & Pasternak 2006; Hussar & Pasternak 2009) and sends task-relevant feedback to IT cortex (Tomita et al. 1999; Kar et al. 2019; Kar & DiCarlo 2021). Given the high level of interconnectivity between visual areas, and feedback projections from frontal areas to visual areas, prior work would suggest that neurons in area IT could possibly access motion information to support object recognition in the face of shape uncertainty.

To test this idea, we developed a behavioral task for monkeys that would make motion behaviorally relevant and necessary for shape recognition. We then recorded neural responses in IT cortex during the task. The task required monkeys to match a shape to a target with variable levels of shape and motion information that could be used to guide responses. On some trials, shape clarity was degraded to a level where only motion information could be used to solve the task. We first used passive viewing stimuli to assess how individual neurons responded to the various levels of target clarity and motion. We then used responses during the active task to look at interactions between target clarity and motion. Interestingly, we discovered that some neurons preferred blurry shapes over crisp ones, and that neurons could fire maximally to a range of shape clarity levels. For one monkey, neurons that responded best to middle-range clarity levels were more likely to exhibit motion sensitivity. For both monkeys, neurons selective for a range of clarity levels also exhibited motion sensitivity.

Reports of IT neurons responding better to degraded images than crisp ones are limited (Namima & Pasupathy 2021), but it is not surprising that degraded images could be coded in this way, as somehow the brain does achieve recognition under various kinds of visual ambiguity (Wyatte et al. 2012; Emadi & Esteky 2013). Most studies of area IT neurons use rapid presentation of clear images in initial surveys for neurons, select cells based on image sensitivity and selectivity using single electrodes, and average across neurons to analyze population responses, all of which could obscure the properties of individual responses (and averaging did obscure individual response characteristics in our data and population-level averages showed no significant differences between task conditions). With the adoption of high channel count recording probes (e.g. NeuroPixel and others), inspection of individual units in real time becomes more and more challenging. This not only necessitates a different kind of neuronal sampling but makes it more likely that one might observe unexpected ranges of selectivity (such as preferred responses for blurred stimuli). Perhaps most neurons in IT cortex respond to clear images, and those that respond better to blurry shapes are sparse. Our data do not support this claim, as a nearly even number of neurons were found having selectivity for blurry shapes as crisp ones, and there appeared to be a gradient of responses (not just two distinct classes). Furthermore, the different kinds of responses could be observed within 100μm of each other on the multi-channel probe, suggesting that neurons with differing response properties are highly intermingled, and possibly interconnected. Future work could investigate the properties of these neurons that appear selective for degraded images. This would require a slightly different approach than most single unit recording studies in IT cortex.

Recent work has shown tuning for blur and occlusion in V4 (Fyall et al. 2017; Oleskiw et al. 2018), which sends direct projections to area IT. It would be interesting and useful to see how neurons in V4 might be transmitting information about stimulus ambiguity to neurons in area IT. Furthermore, this raises the question of whether the motion sensitivity we observed is true motion sensitivity, or “degraded feature” sensitivity. Here we only combined motion with shape, but it is easy to imagine an extension of the task employed here where multiple secondary features (e.g. motion, color, and texture) were bound with shape in a recognition task to determine if there are neurons that support recognition of degraded shapes across all these different feature dimensions (e.g. true multidimensional feature integrators) or whether there are distinct subpopulations responding to degraded features that support shape recognition, and thus object recognition. Such an experiment could also compare different methods of degradation, which might also affect how IT neurons respond to degraded shape stimuli. Perhaps blur is a special kind of degradation of shape, which motion can induce (i.e., motion makes shapes blurry) and thus promotes the interaction we observed in our data at the level of neural responses. More work needs to be done to disentangle motion and blur interactions in area IT, but this study is the first to demonstrate that these interactions exist.

Decoding of neural population activity was also used to demonstrate that neurons in IT cortex could discriminate between motion conditions, even when shape information was not available. We interpret this as a representation of motion. One question that arose from this result was what was being decoded? The motion conditions were decodable in both passive and active tasks, with and without fixation required, with suggesting that eye movements were not leading to the decoding results observed. Furthermore, analysis of motion sensitivity across time showed that the subpopulations of neurons responding to motion changed depending on shape clarity, and that motion sensitivity changed with time, suggesting that trajectory representation is happening at a population level, not in single neurons. As motion trajectories are positional sequences, future work could investigate the distinction between motions that are predictive and non-predictive and leverage previous work on the responses of neurons in area IT to sequential information (Meyer & Olson 2011; Ramachandran et al. 2017; Kumar et al. 2017; Kaposvari et al. 2018). Simultaneous recordings from large groups of IT neurons would also enable analyses to determine how trajectories might be coded within subpopulations of IT neurons; the work presented here was limited to using pseudo-populations as a proxy for widespread joint population signals. This would shed light on what was truly being decoded, whether it was indeed any motion information, or motion information that had been behaviorally linked to shapes (or shape categories).

If area IT has access to motion information as our data suggests, how is this related to general motion responsiveness elsewhere in the STS? A direct comparison with neural responses in STS would be useful for understanding the respective contributions of area IT and STS to motion representation during degraded shape recognition. STS is well known for its selectivity for motion and form and conjunction of these features at the single neuron level (Saito et al. 1986; Oram & Perrett 1996; Jellema et al. 2004; Schultz et al. 2005; Hein & Knight 2008; Jastorff et al. 2012). Recording in STS and area IT simultaneously could provide a reference for gaining a better understanding of what was being decoded from neurons in IT cortex, as well as the different roles that area IT and STS play in moving object recognition.

The data from this experiment cannot be used to discern whether the neurons that respond to motion information when shape is degraded are only motion sensitive, or if they are sensitive to other secondary features and context that might support object recognition when shape information is weak or unavailable. Although such responses have not been commonly reported, it is not surprising that some neurons in area IT respond better to degraded images than crisp ones, as the brain must be capable of somehow recognizing objects under degraded conditions, with limited information, and it is well understood that area IT plays a critical role in object recognition (Conway 2018). A recent study demonstrated that neurons in area IT indeed play a role in recognition of degraded patterns, and although onset latencies were not affected by the level of noise, neurons could be characterized as accumulating evidence (Kuboki et al. 2016), which implies that these neurons must integrate information over time, a critical feature required for coding of motion information. Furthermore, recent work has also demonstrated that neurons in area IT can also encode category-orthogonal properties, including those that are often considered lower-level features, when behaviorally relevant (Hong et al. 2016). Thus, we hypothesize that the neurons we found to be motion sensitive are likely to be sensitive to other features as well, but perhaps play a larger role in population responses in area IT when the primary feature of the area, shape, is degraded or unavailable. This interpretation raises the question of whether such response properties (i.e. responding to features other than those traditionally coded in area IT) associated through task training are present in this region before any learning has taken place? It might be possible to determine this using a task requiring the monkeys to learn to recognize new degraded objects while recording from the same neurons during learning.

In summary, the data here suggest that neurons in IT cortex have access to motion information under certain task conditions. Furthermore, the data show an interaction between sensitivity to motion and sensitivity to shape clarity at the single neuron level, which suggests that motion sensitivity in area IT might have been overlooked due to the use of paradigms using rapid presentation of crisp visual stimuli. The paradigm and data here provide a useful platform for future investigation of motion and ambiguity in area IT, both of which are crucial to a clear understanding of object recognition more generally.

## Acknowledgements

We would like to thank Dr. Theresa Desrochers, Dr. Thomas Serre, Dr. Dima Amso, Dr. Rufin Vogels and Dr. Wilson Truccolo for their comments on early versions of this manuscript and intellectual contributions during this research. We would also like to thank Dr. Shushruth, Dr. Gideon Caplovitz and Dr. Ryan Mruczek for their constructive feedback on drafts of this manuscript. Thank you also to Dr. Shaobo Guan, Dr. Ruobing Xia, Dr. Ryan Miller, Dr. Wenhao Dang, Aarit Ahuja, John Ghenne, and Nadira Yusif-Rodriguez for their various suggestions and insights throughout the course of this work.

